# Mitotic bookmarking by CTCF controls selected genes during the fast post-mitotic genome reactivation of ES cells

**DOI:** 10.1101/2022.02.20.481195

**Authors:** Almira Chervova, Nicola Festuccia, Agnes Dubois, Pablo Navarro

## Abstract

Mitosis leads to a global downregulation of transcription that then needs to be efficiently restored. In somatic cells, this is mediated by a transient hyper-active state that first leads to the reactivation of genes necessary to rebuild the interphasic cell and then of those executing specific cell functions. Here, we hypothesized that cells displaying rapid cell cycles may display accelerated gene reactivation dynamics. To test this, we focused on mouse Embryonic Stem (ES) cells, which have a short cell cycle and spend a minor time in G1. Compared to previous studies, we observed a uniquely fast global reactivation, which displays little specificity towards housekeeping versus cell identity genes. Such lack of specificity may enable the restoration of the entirety of regulatory functions before the onset of DNA replication. Genes displaying the fastest reactivation dynamics are associated with binding of CTCF, a transcription factor that largely maintains binding to its targets on DNA during mitosis. Nevertheless, we show that the post-mitotic global burst is robust and largely insensitive to CTCF depletion. There are, however, around 350 genes that respond to CTCF depletion rapidly after mitotic exit. Remarkably, these are characterised by promoter-proximal mitotic bookmarking by CTCF. We propose that the structure of the cell cycle imposes distinct constrains to post-mitotic gene reactivation dynamics in different cell types, via mechanisms that are yet to be identified but that can be modulated by mitotic bookmarking factors.

## Introduction

The duration of the cell cycle is highly variable between cell types and during developmental and physiological stages^**1**^. A significant amount of this variability depends on the length of the G1 phase^**2**^, the period following mitosis during which the cell prepares for replicating its DNA and undergoing, or not, an additional division. This transition, known as the G1/S checkpoint, is controlled by extrinsic signals, such as growth factors, which enable progression through the remaining phases of the cell cycle when the environment is favourable for cell growth^**3**^. Moreover, the gap between mitosis and replication also provides time for the cell to reinstate an appropriate gene transcription profile. Indeed, mitosis is accompanied by major molecular modifications that entail a drastic downregulation of transcription and a halt of gene regulatory processes^**4**^. Hence, following mitosis, the cell needs to reactivate its regulatory network and gene expression to ensure housekeeping functions such as DNA replication or protein synthesis, and to maintain the specific molecular and physiological features that define its identity^**4**^. How the genome undergoes this reawakening of transcription is not entirely understood. Recently, several genome-wide approaches have addressed this question in somatic cells^**5-7**^. A picture has emerged that a strong burst of transcription characterises around half of active genes, which may reach after mitosis the highest levels of transcription they will ever attain during the cell cycle^**5**^. Such hyper-activation is observed first for genes required to rebuild the interphasic cell, and only later for those linked to cell identity^**6,7**^. In the examples studied with sufficient time resolution after release from a mitotic block^**5,7**^, the vast majority of genes reach maximal transcription levels during a window of 80 to 300 minutes, with the first signs of transcription apparent after approximately 60. Such kinetics are relatively rapid, with respect to the long cell cycles of most cell types, which typically last 24h. Yet, the timely reactivation of the genome might represent a challenge to rapidly proliferating cells. In these cell types, a short G1 phase of less than 2-3h might require replication to start even if transcription is not fully re-established. Alternatively, fast cycling cells might display accelerated reactivation dynamics. In support, transformed cells^**6,7**^ appear to reawaken their genome slightly faster than non-transformed ones^**5**^, suggesting that intrinsic cellular differences influence the pace of post-mitotic events. To understand if this relationship applies also to untransformed cells, we focused on undifferentiated mouse Embryonic Stem (ES) cells. Derived from early embryos, these pluripotent cells display a number of properties that may make fast reactivation dynamics after mitosis essential^**8,9**^. First, their average cell cycle time is of only 12h, due to the absence of a G1/S checkpoint; almost immediately after mitosis ES cells start DNA replication. Second, they self-renew indefinitely, indicating that they are particularly efficient in maintaining their transcriptome across virtually infinite cell divisions. Third, the preservation of their properties strictly relies on a nearly-continuous regulation of gene expression by transcription factors (TFs). Fourth, ES cells were shown to be particularly prone to differentiate after mitosis, indicating that their identity is unstable in G1. We observe that most genes, around 90%, reach maximal transcription levels within the first two hours after division and a substantial fraction initiate their reactivation as soon as 20-30 minutes after release from a mitotic block. These kinetics allow the near complete re-establishment of the transcriptome before the onset of replication. Given that the TF CTCF^**10**^ remains bound to a large fraction of its target sites during mitosis^**11**^, a process that is known as mitotic bookmarking^**4**^ and that is either strongly reduced or completely abrogated in other cell types^**11-13**^, we hypothesized that CTCF may contribute to the particularly fast and global transcriptional bursting occurring in post-mitotic ES cells. Using auxin-inducible depletion of CTCF^**14**^ we rule out such global function for CTCF, highlighting the robustness of the post-mitotic burst in ES cells. Despite this, however, we identify around 350 genes whose appropriate reactivation is tightly regulated by promoter-proximal mitotic binding of CTCF, providing a direct experimental evidence of the mitotic bookmarking function of CTCF. We propose that while yet to identify mechanisms may underlie important cell type specificities of the post-mitotic hyper-active state, mitotic bookmarking factors play an important role in modulating its dynamics and amplitude for selected groups of target genes.

## Results

### Following post-mitotic transcription in ES cells with high temporal resolution

We established experimental conditions to follow gene reactivation after mitosis using CCNA-GFP cell cycle reporters^**15**^ and a double inhibition protocol that allows the efficient block of ES cells in metaphase (4h CDK1 inhibition with RO-3306 followed by a release into nocodazole-supplemented medium for 4h and shake-OFF; **Fig.1A**). We monitored the dynamics of GFP expression and changes in DNA content by Fluorescent-Activated Cell Sorting (FACS), as mitotic cells completed division and re-entered interphase. Two hours post-release, we observed that a substantial fraction of cells had repopulated the G1 compartment (**Fig.1A**). Moreover, we also observed a gradual decompaction of the chromatin and the acquisition of an interphase-like nuclear morphology, characterised by prominent chromocenters, during the first two hours of release (**Fig.1B**). Therefore, we aimed at analysing gene reactivation during this period. Other studies have used direct methods to monitor post-mitotic transcription^**5-7,16**^, such as RNAPII profiling or labelling and purification of nascent transcripts in FACS-purified G1 cells; however, these strategies practically preclude highly resolutive temporal analyses as they require extensive amounts of starting material or manipulation. Hence, in order to generate gene reactivation profiles with high temporal resolution, we decided to use ribo-depleted, single-stranded RNA-seq and a STAR-RSEM pipeline^**17,18**^ that allowed to quantify intron-containing pre-mRNA isoforms. Asynchronous cells were harvested in parallel with mitotic cells and samples obtained at different times following nocodazole release (20, 30, 40, 50, 60, 90 and 120 min) and immediately processed for RNA extraction. Therefore, in comparison to previous studies in ES cells that relied on a maximum of two time-points after mitosis^**15,16,19,20**^, our strategy provides an unprecedented temporal resolution, revealing a previously overlooked complexity. For instance, based on an early (1h post-mitosis) versus late-G1 (3h) analysis, pluripotency-associated genes have previously been shown to reactivate fast and remain highly transcribed^**16**^. However, our time-resolved analysis shows a variety of behaviours within the first two hours of release (**Fig.S1**). Genes such as *Oct4* and *Klf4* display a strong burst at slightly different timepoints, followed by a downregulation; others, like *Esrrb*, reactivate later but remain stably transcribed (**Fig.1C**). Similarly, cell-cycle associated genes have been previously found to reactivate early or late after mitosis, depending on their function during the G1/S or G2/M transition^**16**^. Instead, our results show a large variability in gene transcription patterns (**Fig.S1**), with many cell-cycle genes showing intense, non-synchronous early bursts of transcription that may, or may not, be subsequently maintained (**Fig.S1** and **Fig.1C**). Therefore, without the required temporal resolution, the differential reactivation kinetics characterising individual genes remains unnoticed (**Fig.1C**). As a final control of the quality of our dataset, we monitored transcription of histone genes, which is known to occur specifically during the S phase^**21**^. Accordingly, and in contrast to most other genes (**Fig.S1**), histones displayed much higher transcriptional levels in asynchronous cells than after division, reflecting a prevalence of cells in S phase in cycling ES cells (60 to 70%). Nevertheless, all histone genes showed signs of substantial transcriptional reactivation as soon as 50’ after release (**Fig.1C**), indicating a fast entry into S phase. Hence, previous studies using 1h as an earliest time point after mitosis may have missed the most immediate post-mitotic dynamics, which, presumably, are those tightly linked to processes such as mitotic bookmarking. Thus, our approach captures the kinetics of transcription reactivation after mitosis, and before the onset of the S phase, with unprecedented details.

**Fig. 1.**
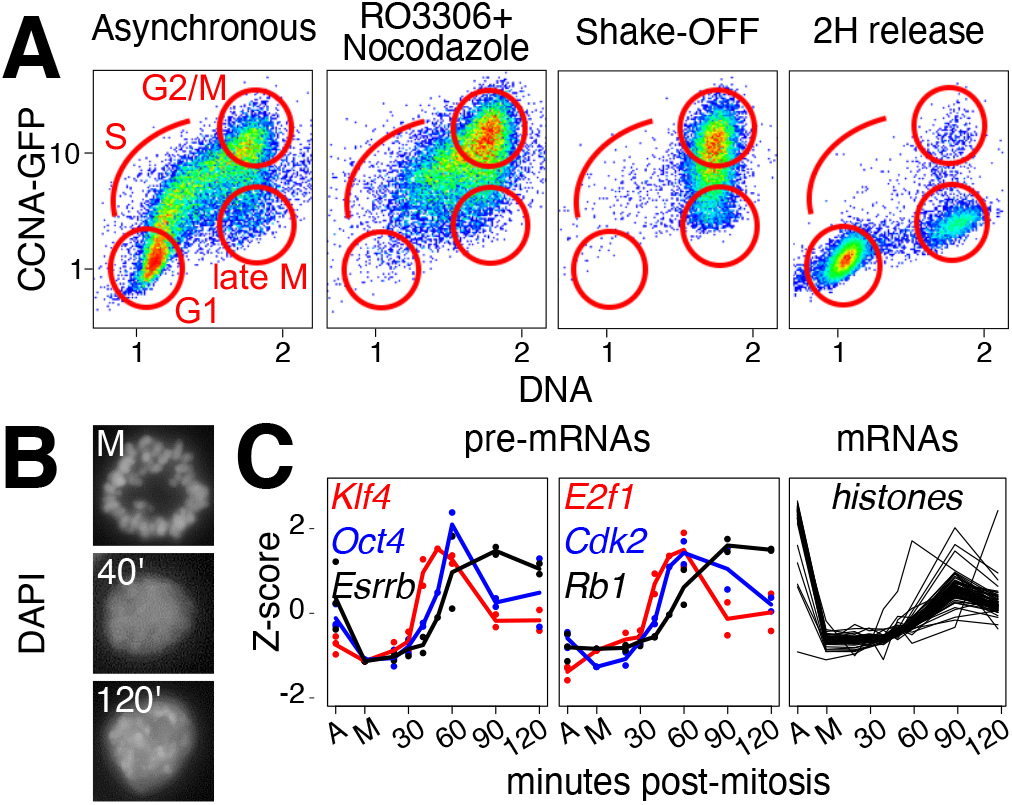
Following gene reactivation in post-mitotic ES cells with high temporal resolution. **(A)** FACS scatter plots of CCNA-GFP cell-cycle reporters (Y-axis, CCNA-GFP fluorescence, arbitrary units, log scale; X-axis, relative DNA content measured by Hoechst staining). Four conditions are presented, as shown above each panel. Cell cycle phases are indicated in red in the plots. **(B)** Representative image of cells stained with DAPI upon mitotic arrest and shake-off (top), and after 40 and 120 minutes post-release (middle and bottom). Note obvious chromocenters (DAPI-dense regions) after 120 minutes, indicating the restoration of a chromatin organisation typical of interphase. **(C)** Examples of post-mitotic gene reactivation (Y-axis, z-score; X-axis, timepoints. A indicates asynchronous cells, M mitotic cells and the numbers represent the minutes after release from the mitotic block). Dots show individual replicates and lines averages (n=3 for A and M; n=2 for each post-mitotic sample). Pre-mRNA levels are shown for pluripotency genes (*Oct4, Klf4, Esrrb*) and cell cycle regulators (*E2f1, Cdk2, Rb*); mRNA levels for individual histone genes (only averages are shown).

### Rapid genome-wide transcriptional bursting in postmitotic ES cells

We then aimed at studying the global dynamics of gene reactivation. For this, we focused on around 10,000 Refseq curated genes displaying at least 0.5 pre-mRNA Transcripts Per Million (TPM) in either asynchronous cells, or in 2 or more samples after nocodazole release (**Table S1**). These were then ranked according to the mean post-mitosis fold-change between 20 and 90’ of release. We observed a fast and global reactivation of the transcriptome, starting 20 to 40’ for most genes, reaching particularly high levels within an hour and declining thereafter (**Fig.2A**). This intense burst of transcriptional activity has been previously observed^**5-7**^, even though in ES cells it occurs faster and the magnitude and the number of genes engaged appear to be substantially higher. Indeed, 90% of genes reach their maximal transcriptional level earlier than 2h following mitosis (**Fig.2B**, left) and around 30, 65 and 80% reached or surpassed the levels measured in asynchronous cells at 40, 60 and 120’, respectively (**Fig.2B**, right). This analysis also identified around 10% of genes that displayed low levels in asynchronous cells but a strong burst after mitosis, possibly reflecting genes with preferential transcription during G1. Principal component analysis (PCA; **Fig.2C**) confirmed that major global changes take place between 40 to 60’ after mitosis. Following this, the transcriptome became increasingly similar to that of asynchronous cells. Notably, the most pronounced differences to asynchronous cells were observed after 50-60’, coinciding with the maximal transcriptional peak. PC1 and PC2 capture 80% and 12% of the measured variance, respectively. Therefore, we selected the top and bottom 1000 genes contributing to PC1 and PC2 (PC1t, PC1b, PC2t, PC2b in **Fig.2D**) to explore 4 major dynamics contributing to the temporal changes of the transcriptome (**Fig.2D**): PC1t and PC2t genes are detected at high levels in asynchronous cells and are either rapidly (PC1t, in red in **Fig.2D**) or more slowly reactivated after mitosis (PC2t, in orange in **Fig.2D**); PC1b and PC2b genes are detected at low levels in asynchronous cells and either remain poorly transcribed (PC1b, in black in **Fig.2D**) or display a strong and transient transcriptional burst after mitosis (PC2b, in blue in **Fig.2D**). Gene Ontology analyses with these 4 groups of 1000 genes characterised by different transcriptional behaviours revealed that genes contributing to PC1t are prominently enriched in global cellular functions, the G1/S cell cycle transition and replication, but strikingly also in stem cell related terms including the pluripotency network (**Fig.S2**). Selection of all significant associations (FDR<0.05) and calculation of the FDR corresponding to their enrichment along a sliding window of 500 genes displaying increasingly fast reactivation dynamics (from top to bottom in **Fig.2A**), showed a strong bias towards the fastest genes (**Fig.2E**). GO terms included global gene expression functions, the G1/S transition and DNA replication, as well as the pluripotency network. Altogether, we conclude that ES cells display less functional specificity regarding how the genome awakens after mitosis, compared to other cell types where genes associated with the rebuilding of the interphasic cell precede cell type-specific genes^**6,7**^. The nearly simultaneous and fast reactivation of the cell cycle machinery, basal gene expression functions and the pluripotency network in ES cells, may enable these cells to cope with a short cell cycle, with the lack of a G1/S checkpoint, and satisfy their continued dependency on TFs to self-renew^**8**^.

**Fig. 2.**
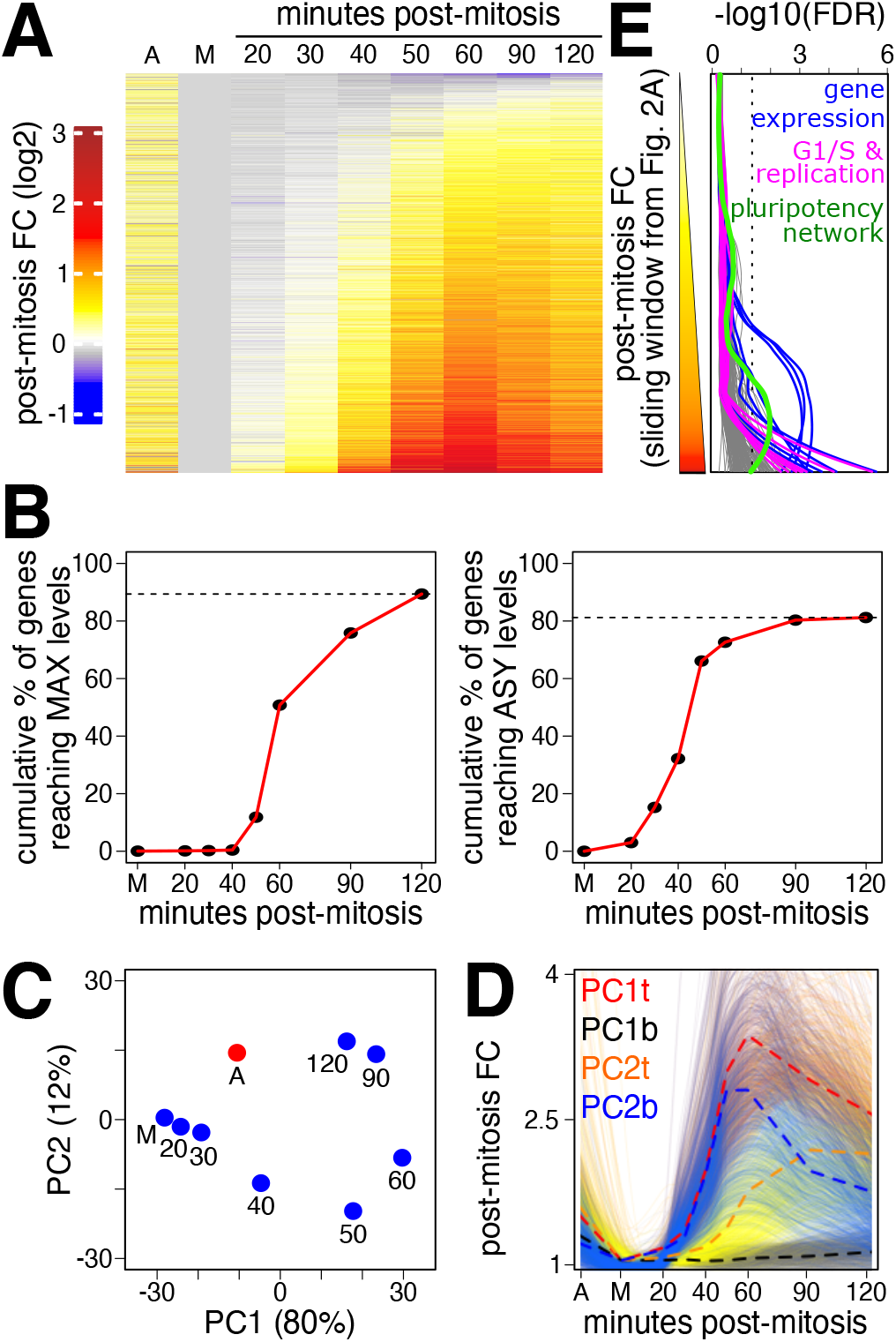
A prominent and fast burst of transcription shortly after mitosis. **(A)** Heatmap showing average log2 fold change to mitosis of pre-mRNA levels for 9911 individual genes in asynchronous cells (A), mitotic cells (M), and at each timepoint post release, ranked from top to bottom by the mean of 20 to 90 minutes. **(B)** Cumulative plots showing the percentage of genes reaching maximal detection levels (left) or the levels detected in asynchronous cells (right) as a function of time. Note that around 10% of genes (1033) display maximal transcription in asynchronous cells. An additional 10% of genes (831) showed instead low levels of transcription in asynchronous cells compared to mitosis. **(C)** Principal Component Analysis of all considered samples. The variance captured by each dimension is indicated. Samples are labelled as in Fig.2A. **(D)** Individual traces (light lines; smoothed with a loess regression) and mean profiles (dashed bold lines) of 1000 genes identified from the top and bottom PCA loadings of PC1 and PC2, revealing the major tendencies driving post-mitotic gene reactivation (Y-axis, fold change to mitosis; X-axis as in Fig.1C). These 4 gene lists were subject to gene Ontology analyses, as shown in Fig.S2. **(E)** Plot showing the -log10(FDR) for the enrichment of a variety of gene ontology terms in groups of 500 genes, defined by sliding a window over genes ranked based on their reactivation kinetics, as in Fig.2A (top, weak post-mitotic reactivation; bottom, strong post-mitotic reactivation). The ontology terms plotted are those enriched in the four groups of genes defined in Fig.2D. Note that all categories of terms show increased enrichment for gene groups displaying strong reactivation dynamics after mitosis. Illustrative categories are coloured: Gene Expression in blue – mRNA processing (GO:0006397), mRNA splicing (GO:0000398 and GO:0000377), gene expression (GO:0010467), ribosome biogenesis (GO:0042254), translation (GO:0006412), among others –, the G1/S transition and replication in magenta – G1/S transition of mitotic cell cycle (GO:0000082), DNA-dependent DNA replication (GO:0006261), pre-replicative complex assembly (GO:0036388), DNA replication initiation (GO:0006270), among others, and the Pluripotency Network in green as defined in WikiPathways (PluriNetWork WP1763).

### Proximal CTCF binding correlates with post-mitotic gene transcription

We next aimed at exploring whether gene reactivation after mitosis was correlated with the binding of specific TFs, which are the main drivers of ES cell self-renewal and pluripotency^**8**^. To do this, we selected 4 groups of genes reaching or surpassing the levels detected in asynchronous cells (as in **Fig.2B**, right panel) at 40 (n=3188 genes), 50 (n=3362), 60 (n=647) or 90 (n=762) minutes post-mitosis, a fifth group (labelled late in **Fig.3A,B**; n=1121) that never reached the levels detected in asynchronous cells and a sixth group showing low levels in asynchronous cells but a marked increased after mitosis (labelled G1_high in **Fig.3A,B**; n=831). Next, we computed the percentage of genes of each group bound by pluripotency TFs (Nanog^**22**^, Oct4^**19**^, Sox2^**19**^ and Esrrb^**19**^) or by the architectural TF CTCF^**10,14**^ within a 25kb region centred on the transcription start site. Strikingly, only CTCF clearly showed increased binding near genes reactivating fast (**Fig.3C**; other genomic sizes than 25kb gave similar results, **Fig.S3**). Since CTCF has been shown to display robust mitotic binding at a large subset of its interphase targets in ES cells^**11**^, we further distinguished between genes associated with CTCF during mitosis or only during interphase (bookmarked and lost, respectively, in **Fig.3D**). We found a higher enrichment of bookmarked compared to lost sites for both the most rapidly reactivated (t40) and the G1_high groups. Moreover, when we counted the number of distinct TF binding sites within 25kb centred on each promoter (**Fig.3E**), we found a similar distribution among reactivation groups for all TFs except for CTCF. An increased proportion of genes in the fast reactivating groups (t40, t50 and G1_high groups) associated with 3 or more binding sites, an effect that is attributable specifically to bookmarked, and not lost, CTCF sites (**Fig.3E**). Finally, we computed the distribution of CTCF binding levels in interphase and mitosis in proximity of each group of genes (**Fig.3F**) and observed that fast reactivating groups (t40 and G1_high) are partially depleted of regions showing low levels of CTCF binding. Regions characterised by high binding levels in mitosis are instead enriched near these genes. Taken together, these correlative analyses point to CTCF as a regulator of the reactivation dynamics after mitosis. Notably, the potential effects of CTCF binding are particularly prominent for G1genes, indicating a specific function after mitosis for this TF, which can be uncoupled from its activity during interphase.

**Fig. 3.**
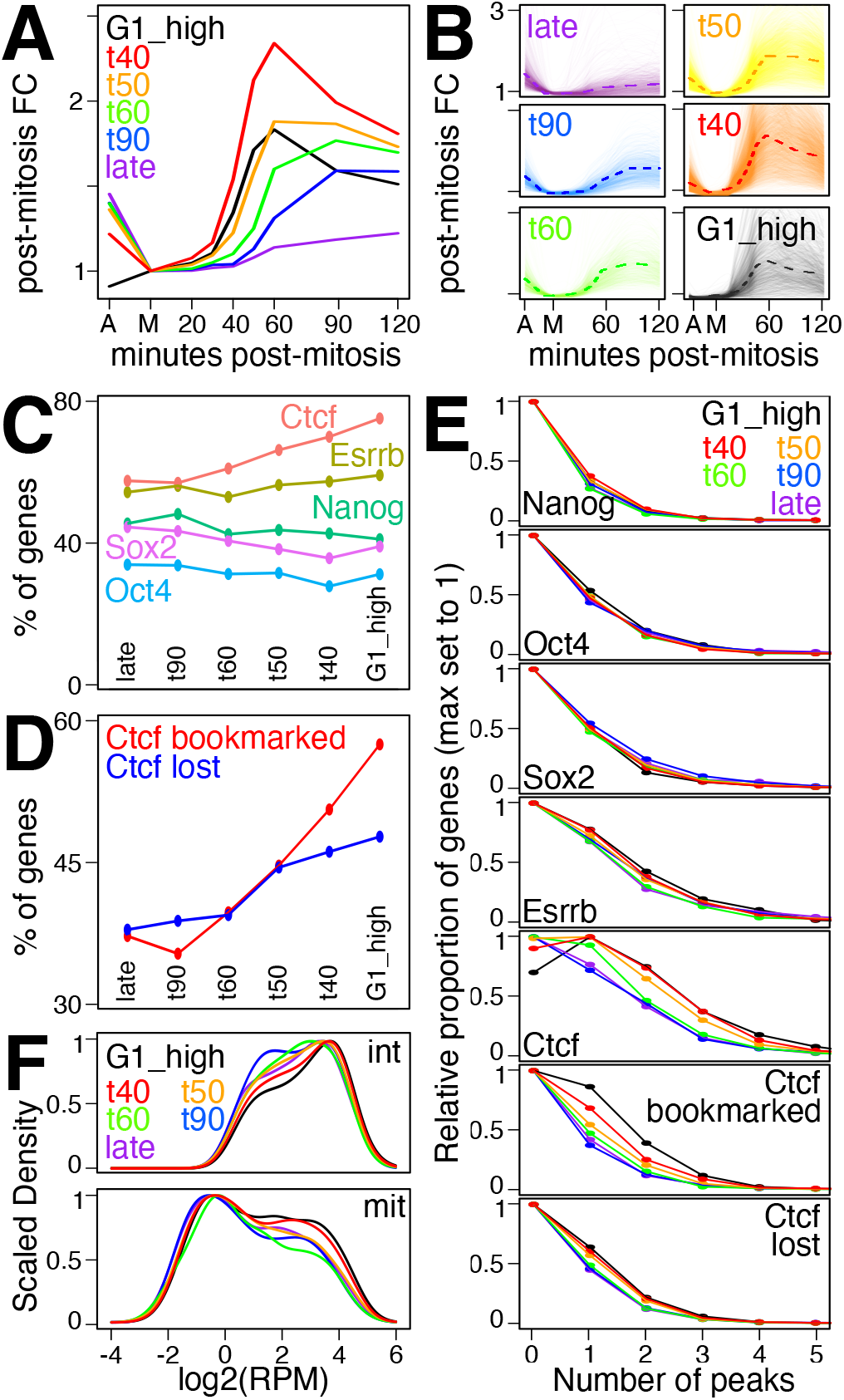
CTCF binding in interphase and mitosis correlates with fast post-mitotic gene reactivation. **(A)** Average post-mitotic fold change of gene groups identified by the time at which the levels detected in asynchronous cells are reached or surpassed, as shown in Fig.2B (G1_high: 831 genes with low detected levels in asynchronous cells; t40: 3188 reaching or surpassing asynchronous levels by 40 minutes post-mitosis; same for t50 to t90: 3362, 647, 762; late: genes reaching asynchronous levels after 120 minutes (88) or never within the analysed post-mitotic timepoints (1033). **(B)** Individual gene traces (light lines; smoothed with a loess regression) corresponding to each category presented in (A), and respective averages (dashed lines). **(C)** Percentage of genes presenting at least 1 TF binding site as detected by ChIP-seq within 25kb centred on the TSS, calculated for each gene category shown in (A). **(D)** Similar analysis to (C) but computing the percentage of genes with mitotically bookmarked and lost CTCF binding sites. Genes were counted as bookmarked or lost when at least one or no CTCF binding site was detected in interphase and mitosis, respectively. **(E)** The relative proportion of genes for each reactivation category shown in (A) as a function of the number of binding sites (X-axis) for the indicated TFs. Binding sites overlapping a 25kb region centred on the TSS were considered. All distributions are scaled to the maximal proportion, observed for zero sites for all TFs, except for CTCF, for which it was at 1 site for rapidly reactivating gene groups. **(F)** Distribution of CTCF binding levels in interphase (top) and in mitosis (bottom) for each gene group shown in (A). For each gene, all the binding sites overlapping a 25kb region centred on the TSS were considered. Binding levels (X-axis) are expressed as reads per million (RPM).

### Robust post-mitotic transcriptional reactivation in the absence of CTCF

To test the role of CTCF in post-mitotic gene reactivation, we took advantage of CTCF-AID ES cells^**14**^, which enable fast (2h) and acute degradation of CTCF upon auxin (IAA) treatment as we previously showed^**11**^ by western-blot, immunofluorescence, ChIP-seq and MNase-seq. CTCF-AID ES cells were synchronised in mitosis as described above, except that they were cultured in the presence/absence of IAA during the last two hours of nocodazole treatment. Subsequently, mitotic cells were shacked-off and re-seeded in nocodazole-free medium -/+ IAA such that they completed mitosis and re-entered interphase with or without CTCF. Despite the global association of gene reactivation with CTCF binding (**Fig.3**), very similar profiles were observed in the presence (-IAA) or absence (+IAA) of CTCF (**Fig.4A** and **Table S2**). Principal Component Analysis (PCA) revealed similar trajectories of the +/-IAA transcriptomes (**Fig.4B**), leading to nearly identical median gene reactivation dynamics (**Fig.4C**). These results indicate that the post-mitotic burst in ES cells is robust and largely insensitive to CTCF depletion, as recently shown in other biological contexts^**12,23**^. Nevertheless, careful examination of the PCA (**Fig.4B**) suggests that small differences exist along PC1 between + and -IAA treatments, 20 and 40min after mitosis. At later time-points, small differences appear instead in PC2. In contrast, both +/-IAA datasets in asynchronous and 2h post-mitosis showed identical PC1 and PC2 scores, providing an internal control to the small but measurable effects mediated by CTCF as cells exit mitosis. Hence, while the global effect of CTCF following mitosis is undeniably small, these observations prompted us to identify gene-specific responses that could be more directly associated with CTCF binding, especially in mitotic cells.

**Fig. 4.**
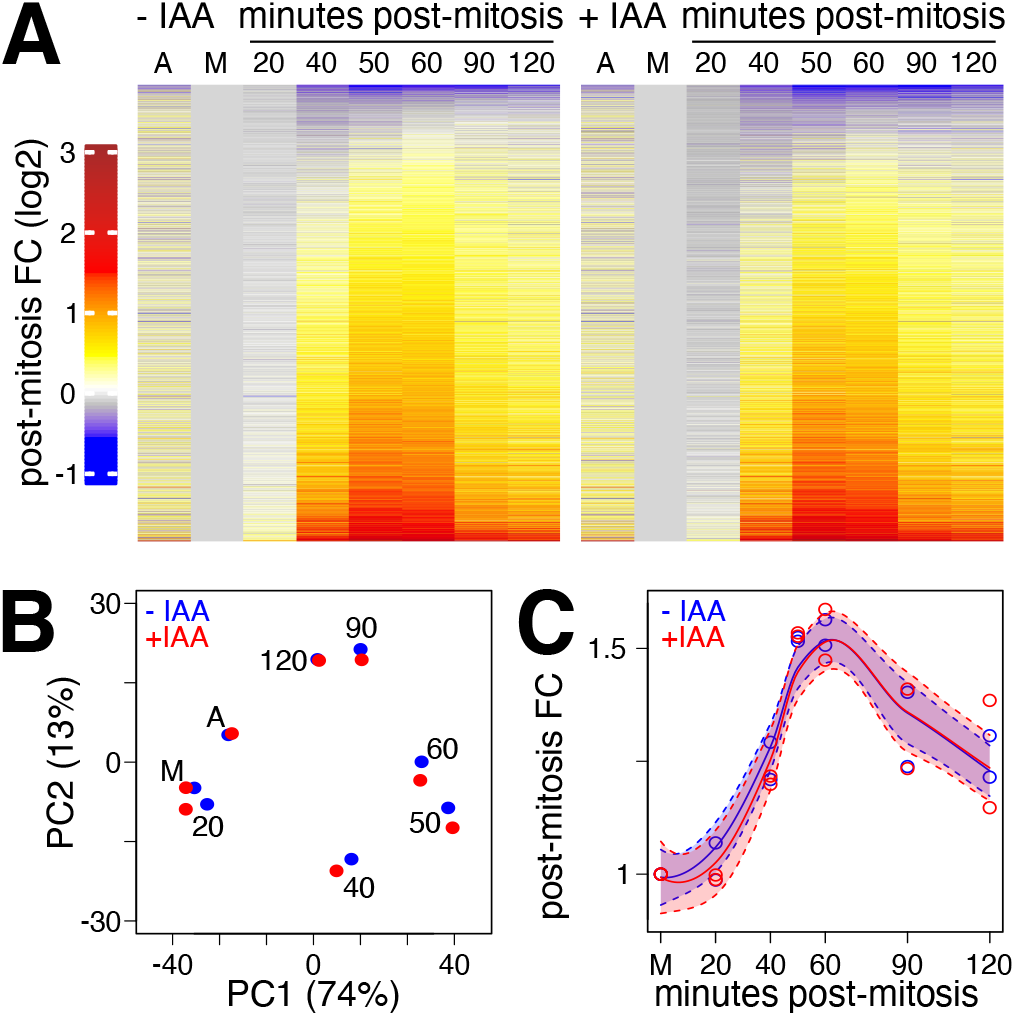
CTCF does not influence global post-mitotic gene reactivation. **(A)** Heatmaps presented like in Fig.2A for untreated (-IAA; left) or CTCF-depleted (+IAA; right) CTCF-AID cells. **(B)** PCA of post-mitotic gene reactivation in the presence (-IAA; blue) or absence of CTCF (+IAA; red). **(C)** Smoothed (loess) median reactivation kinetics in the presence (-IAA; blue) or absence of CTCF (+IAA; red). The shadow areas represent 95% confidence intervals of the regression.

### Mitotic bookmarking of CTCF is required for proper reactivation of selected promoter-proximal target genes

Using standard RNA-seq differential expression analysis (DEseq2^**24**^), we identified 611 genes displaying differential pre-mRNA expression in +/- IAA in either asynchronous or in at least two post-mitosis samples (**Fig.5A** and **Table S2**). These 611 genes were further divided in genes being up- or down-regulated upon the loss of CTCF and by their temporal response: exclusively in asynchronous cells, or also at either late or early time-points post-mitosis (asy, late and early in **Fig.5A**, with examples in Fig.S4). Among all 6 categories, genes activated by CTCF from early post-mitotic time-points onwards (early_down) were prominent (260 genes). Gene ontology analyses, however, did not identify any relevant term being enriched for any of the identified groups (**Fig.S4**). We conclude, therefore, that while at a global scale the loss of CTCF in mitosis and early interphase is largely inconsequential, a sizeable fraction of genes shows a clear CTCF dependency to be efficiently reactivated. Next, we explored whether the different speeds in responding to CTCF after mitosis were associated to CTCF binding events. For this, we computed Fisher exact tests for the enrichment of each group of differentially expressed genes at increasing distances to the nearest bookmarked or mitotically lost CTCF peak (**Fig.5B**). While asy genes were not enriched, suggestive of indirect effects, we observed that genes activated by CTCF late after mitosis (late-down) displayed low enrichment levels, particularly near Lost CTCF binding sites. Hence, at these genes, the reacquisition of CTCF binding after mitosis is accompanied by their sustained activation, indicating that even in the absence of mitotic binding, CTCF may contribute to gene activity before the onset of replication. This interpretation is supported by the fast rebinding of CTCF to DNA following mitosis^**12,25**^. Strikingly, genes responding rapidly after mitosis to the loss of CTCF displayed a strong statistical association to CTCF binding sites, with a very prominent enrichment of genes activated by CTCF (early-down) within 10kb of CTCF Book sites (**Fig.5B**). This indicates that promoter-proximal bookmarked CTCF sites lead to CTCF-dependent transcription inmediately after division. Thus, mitotic bookmarking confers almost immediate post-mitotic responsiveness to CTCF compared to other non-bookmarked but CTCF-dependent genes. To further confirm these statistical correlations we directly assessed CTCF binding in interphase and in mitosis at asy-only, late and fast responders, using available ChIP-seq data^**11**^. Robust binding at the TSS in both interphase and mitosis was observed at early-responders exclusively, whereas Cohesin^**10**^ (Smc1) was similarly associated with the 3 groups (**Fig.5C**). This data strongly suggests that promoter-proximal CTCF binding in mitosis is associated with early activatory function, demonstrating the mitotic bookmarking role of CTCF in mouse ES cells^**11**^. Moreover, a lower but significant enrichment of CTCF Book sites was also observed for genes being rapidly repressed by CTCF (early-up; **Fig.5B**) indicating that mitotic CTCF binding may also be used as a mean for bookmarking for repression.

**Fig. 5.**
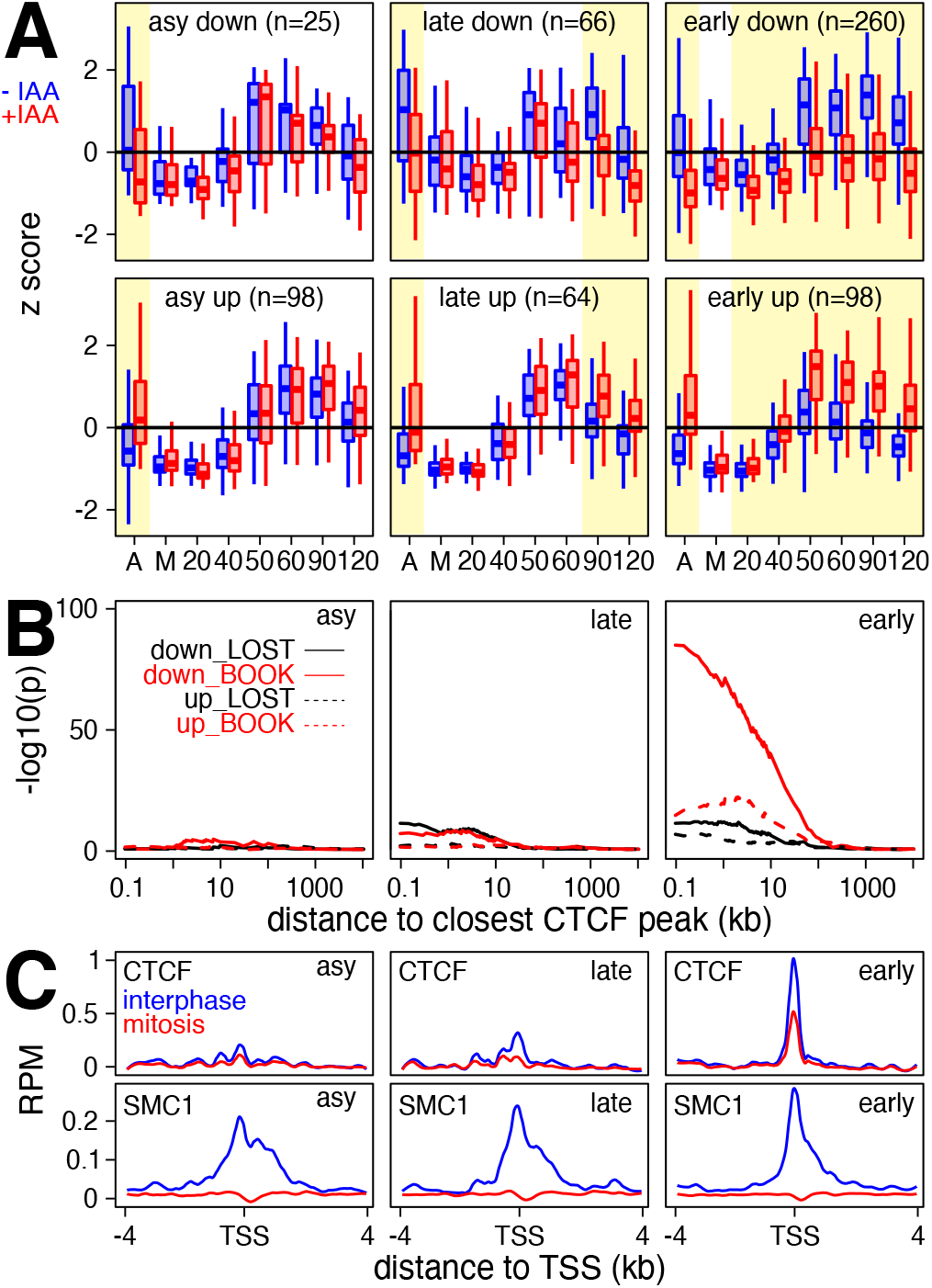
Mitotic bookmarking effects mediated by CTCF. **(A)** Gene expression changes (z-score) in asynchronous cells and during post-mitotic release of CTCFAID cells treated or not with IAA to deplete CTCF (blue, presence of CTCF; red, absence). Samples are indicated on the X-axis. Six gene groups are shown, based on the time of their response to CTCF depletion (asy, late, early) and the direction of the transcriptional change (up, down). The number of genes of each group is indicated. The shadow areas indicate times showing expression changes. **(B)** Statistical enrichment (Y-axis, -log10(Fisher p-value)) of CTCF binding at increasing distances (log10(kb)) of the TSS of genes responding to CTCF depletion exclusively in asynchronous cells (asy; left) or either late (middle) or early (right) after mitotic release. Two categories of CTCF sites are analysed: those that are present in interphase and mitosis (Book, in red) and those that are exclusive of interphase (Lost, in black). The direction of gene responses is also considered (down and up, see (A), as solid and dashed lines respectively). **(C)** Average binding profile (X-axis, reads per million) of CTCF (top) and the Cohesin subunit SMC1 (bottom), across 8kb centred on the TSS of genes responding to CTCF depletion exclusively in asynchronous cells (left) or either late or early upon mitotic release (middle and right, respectively). Interphase is shown in blue and mitosis in red.

## Discussion

Here, we aimed at characterising how ES cells reactivate their genome after mitosis. While several studies have already addressed this question^**15,16,19,20**^, none had the temporal resolution we achieved. We observe that the vast majority of active genes undergoes a prominent transcriptional burst largelly surpassing the levels detected in bulk asynchronous cells. A post-mitotic hyper-active state has now been observed in different cell types^**5-7**^, suggesting that it may represent a general feature. The drivers of this phenomenon remain to be directly identified. Two scenarios can be envisioned. First, the burst observed in G1 could depend on specific and direct mechanisms, such as the behaviour of general and/or ubiquitous TFs executing such function. For instance, the sudden relief of inhibitory phosphorylations on many components of the basal transcriptional machinery might contribute to a synchronous activation of transcription^**26,27**^. Moreover, sequence-specific TFs involved in hyper-transcription in ES cells, but possibly also in other cell types, such as *c-Myc*, could also represent important candidates^**28,29**^. Second, the post-mitotic hyper-active transcriptional state could be the indirect consequence of other properties of post-mitotic cells. For instance, the re-folding of the genome into topologically associating domains (TADs), which largely confine enhancer function, has been shown to take place, in average, after gene reactivation^**12,25,30**^. Importantly, before TADs are reestablished, intra and inter-TAD enhancer-promoter contacts drive inappropriate gene activation^**25**^. Hence, perhaps transiently and randomly established contacts during the transition from a Condensin-mediated mode of 3D organisation in metaphase, to one driven by Cohesin in telophase^**31**^, could fuel the burst of transcription observed in G1. However, a main driver of TADs formation, CTCF^**14**^, does not influence the global dynamics of reactivation, neither in ES cells nor in somatic cells^**12**^. Additional properties, such as a sudden decondensation of the chromatin, should be envisioned as potential drivers. Understanding whether post-mitotic transcriptional hyper-activity is the primary result of specific regulations and/or a secondary effect of other cellular changes will require additional studies.

Compared to published studies performed in other cell types^**5-7**^, ES cells appear to reactivate their genome faster as they exit from mitosis. CTCF fulfils key characteristics that made it as a suitable candidate to explain this particularly expedite reactivation: it displays extensive mitotic binding specifically in ES cells^**11-13**^ and is frequently bound in the vicinity of rapidly reactivated genes. Despite these correlations, our data shows that the post-mitotic burst of transcription occurs in ES cells with overall similar dynamics in the presence or absence of CTCF. Therefore, other mechanisms of post-mitotic acceleration should be investigated. Moreover, our analyses have also shown that the most strongly induced genes after mitosis correspond to genes associated with the G1/S transition and DNA replication, an observation that is in line with the lack of a G1/S checkpoint in ES cells^**8,9**^. Indeed, establishing a functional replication machinery rapidly after mitosis seems a major requirement to efficiently and productively enter into the S phase. However, the temporal differences with other functional categories is minor, below 20 minutes. Therefore, ES cells seem to simultaneously reactivate cell identity and ubiquitous genes, in contrast with other cell types^**5-7**^. This faster and indistinctly global reactivation in post-mitotic ES cells may have evolved as a response to two requirements: one, undergo fast proliferation, achieved by the lack of the G1/S checkpoint^**9**^; two, preserve a continuosly active regulatory network – even subtle changes in TF dosage have been shown to induce the loss of self-renewal and to promote differentiation^**8**^. Hence, it seems reasonable to think that to execute multiple tasks efficiently before entering replication, ES cells may accelerate the global rate of gene reactivation compared to somatic cells that can rely on decision-making mechanisms to transit to the S phase. Nevertheless, caution must be taken as differences between the methods used to synchronise cells or to measure transcriptional outputs may bias, at least partially, the conclusion drawn by direct comparison between studies. However, the fact that ES cells reactivate their transcriptome faster than other cell types is in line with their general hypertranscription^**28**^ and permissive chromatin state^**32**^. Whether these properties are alone sufficient to accelerate post-mitotic gene transcription requires further investigation.

Despite not having global effects, CTCF exerts, however, a clear influence on around 350 genes where it binds at or nearby the promoter, both in interphase and in mitosis. For more than 250 of these genes, mitotic bookmarking by CTCF is required for proper reactivation as soon as 20 min post-release; for around 100, it is conversely required to attenuate transcription early in G1. While it remains unclear how mitotic CTCF binding conveys either activatory or repressive function, these observations experimentally establish that CTCF acts as a mitotic bookmarking factor, as previously suggested^**11**^. Hence, while its main function is generally considered that of shaping the global architecture of the genome, this study underscores the function of CTCF as a gene-specific regulator displaying a key role in the control of selected genes as cells exit mitosis. Whether other mitotic bookmarking factors display such function and are capable of modulating positively or negatively the post-mitotic burst remains to be directly assessed. Moreover, this study calls for reconsidering, at least partially, the role of mitotic bookmarking^**4**^. Indeed, bookmarking by TFs has been long considered as a potential mechanism to accelerate gene reactivation after division. However, the fast and efficient reactivation of the whole transcriptome – particularly in ES cells but also in other cell types – raises the question of the real advantage conferred by this phenomenon. In other words, does anticipating the resumption of gene expression by few minutes, and for a relatively small subset of genes, play a determinant role in preserving cell function? Perhaps counter-intuitively, our data argues against the simple idea of mitotic bookmarking as an accelerator of gene reactivation: CTCF-bookmarked and responsive genes do not reactivate faster, in average, than non-bookmarked genes. Instead, we observe that bookmarked genes are responsive to CTCF early after mitosis, while non-bookmarked genes are much less prone to CTCF-dependent regulation, even if both sets respond to CTCF in asynchronous cells. Moreover, the main effect of mitotic bookmarking does not seem to be changing the timing but, rather, the amplitude of gene reactivation. Therefore, our data suggests that mitotic bookmarking confers early responsiveness rather than rapid reactivation. Whether this is a specificity of CTCF or a general and unapreciated intrinsic feature of mitotic bookmarking by TFs, will be an important question for future studies.

## Methods

### Cell culture and mitotic preparations

ES cells (CCNAGFP^**15**^ and CTCF-AID^**14**^ clone EN52.9.1) were cultured on standard FCS/LIF conditions^**11**^. To obtain mitotic ES cells (>95% purity as assessed by DAPI staining and microscopy), we first arrested them in G2 with 10 µM of CDK1 inhibitor RO-3306 for 4h (Sigma, SML0569), followed by release into 50 ng/ml nocodazole (Sigma, M1404; 4h). Cells were washed in PBS, and a gentle shake-off was then performed in 10 ml of PBS, monitoring the process under a microscope to avoid detaching clumps of interphase cells. The cell suspension was filtered (10 µm filter; pluriSelect, 4350010-03) by gravity and cells spun down, resuspended in pre-warmed medium and immediately transferred to the incubator in pre-warmed 3 cm dishes, that were intentionally not coated with gelatine. For each timepoint/treatment, 1-2 × 10^6^ cells were seeded in separate dishes in a volume of 1 ml, so that upon collection only the cells to be harvested were taken out of the incubator. Cells were detached from the dish by pipetting, spun down at 4°C (1 min, 150 g in a benchtop centrifuge), and lysed in 500 µl of cold TRIzol (ThermoFisher, 15596026). The procedure was also monitored by FACS analysis of CCNA-GFP, cultured in the presence of 20 µM Hoechst-33342 for 20 min, trypsinised, filtered through a 40 µm cell strainer and kept on ice. Hoechst and GFP levels were analysed using a LSR Fortessa instrument (Becton-Dickinson) and the FlowJo software (Tree Star). CTCF depletion in CTCF-AID cells was achieved with 0.5 mM auxin (IAA Sigma, I5148) for 2h, which leads to an acute depletion^**11**^, starting after 2h of nocodazole treatment. IAA was maintained throughout the whole release procedure. Asynchronous cells were treated in parallel during 4h. For each condition, we generated 3 independent replicates. All samples of one replicate were sequenced; for the other two, asynchronous and mitotic cells were sequenced together with alternating timepoints during the release (**Tables S1,S2**).

### RNA-seq

RNA was extracted with 500 µl TRIzol (ThermoFisher, 15596026) and treated with DNAse I (Qiagen, 79254) for 20min at 37°C. Following phenol:chloroform purification, ribo-depleted, stranded and paired-end libraries were prepared and sequenced by Novogene Co Ltd. Reads were aligned to the mm10 genome using STAR^**17**^, quantified by RSEM^**18**^ and counts transformed into Transcripts per Million (TPM, Tables S1 and S2). Gencode (vM12) gtf was used after adding a pre-mRNA isoform for each annotated gene, spanning the longest isoform. Raw data is available at Gene Expression Omnibus (GSE196889).

### Bioinformatic analyses

Analyses were performed in R (3.6.3). Genes with mean TPM>0.5 in asynchronous or in >2 samples post mitosis and present in the Refseq curated repository ([UCSC](https://genome.ucsc.edu/cgi-bin/hgTables)) were considered. Post-mitotic fold-changes were calculated per replicate and then averaged to generate gene expression plots and heatmaps (Complex Heatmap package^**33**^). Smoothed profiles used for visualisation purposes only were obtained with a loess regression. PCA was run with prcomp function with centred data corresponding to post-mitotic log2(FC) for CCNA-GFP and log2(TPM) for CTCF-AID to capture direct differences between IAA treatments. GO analyses were performed with enrichR^**34**^. To correlate genes and TF binding and to compute CTCF/Smc1 enrichments, we first selected a single TSS for each gene, which corresponded to the TSS displaying highest levels of RNAPII, H3K4me3 and DNaseI accessibility using available ES cell data from the Encode consortium. TF binding sites were previously reported^**11,19,22**^. Fisher-exact tests to assess gene-TF proximity enrichments were performed as previously described^**11,19,22,35**^. ChIP-seq enrichments were calculated with [bamsignals](https://bioconductor.org/packages/devel/bioc/vignettes/bamsignals/inst/doc/bamsignals.html). Differentially expressed genes were identified using rounded RSEM counts and DESeq2^**24**^. Pre-mRNAs with FDR<0.1 and absolute log2FC>0.2 in either asynchronous cells or in at least two post-mitotic samples were considered as differentially expressed.

## Acknowledgements

The authors acknowledge E Nora and B Bruneau for CTCF-AID ES cells; N Owens for discussions on experimental design and bioinformatic analyses and J Dekker and M Oomen for discussions; M Cohen-Tannoudji, T Gregor, G Cecere and G Blobel for critical reading of the manuscript. PN acknowledges the Institut Pasteur, the CNRS, and Revive (Investissement d’Avenir; ANR-10-LABX-73) for recurrent funding and the Ligue contre le Cancer (LNCC EL2018 NAVARRO) and the European Research Council (ERC-CoG-2017 BIND) for financial support.

## Author contributions

NF generated the data with help from AD; AC made bioinformatic analyses. PN conceived the project, analysed the data and wrote the manuscript with NF.

## Supplementary information

Four supplementary figures accompany this manuscript, they can be found at the end of this document. Two Supplementary Tables are available online.

---

**Supplementary Information, Fig.S 1.**
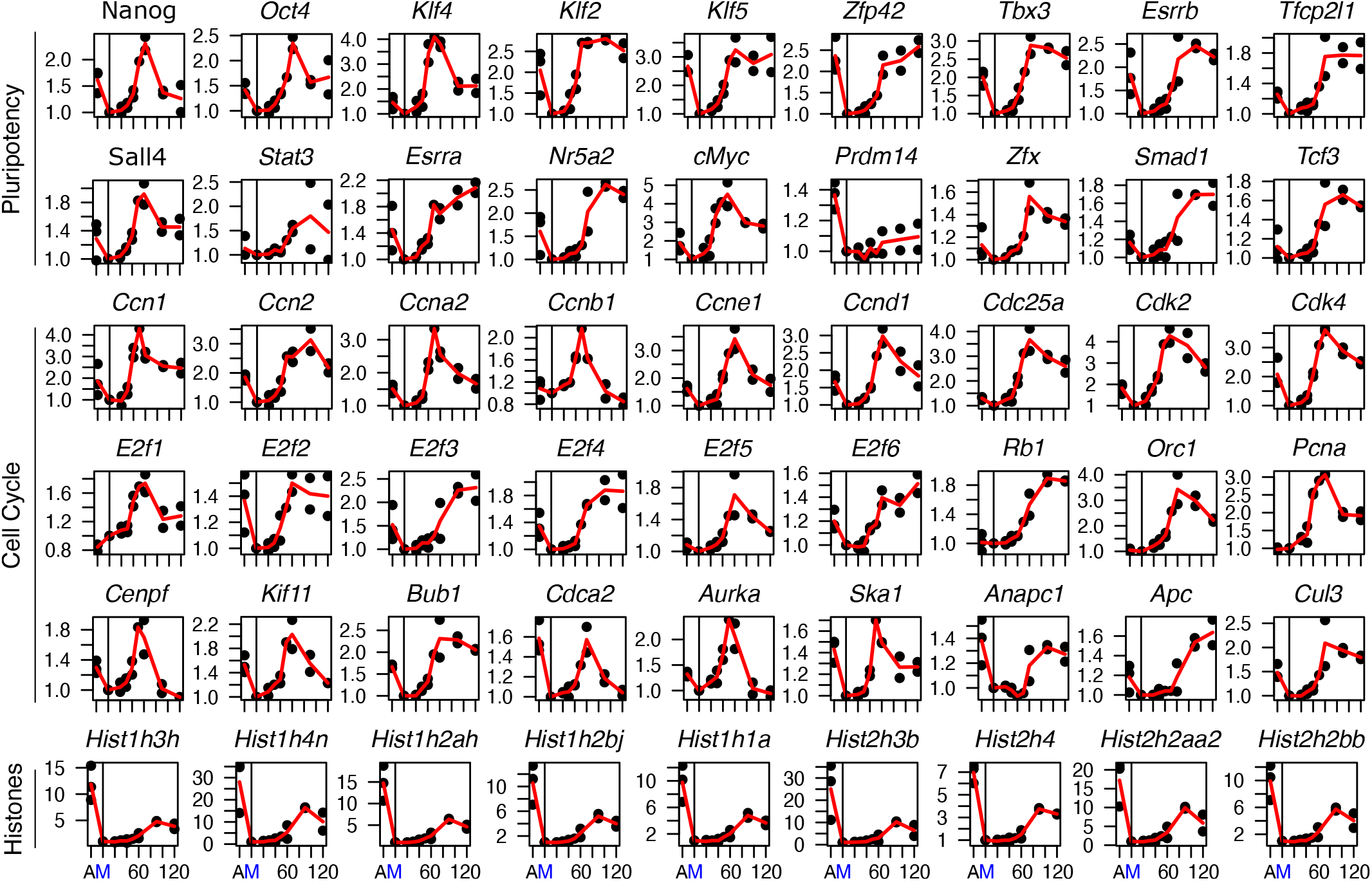
Examples of transcriptional profiles for individual genes. For each plot, each dot represents a replicate and the red line the corresponding average. Y-axis shows fold change to mitosis; X-axis the samples (A: asynchronous cells; M: mitotic cells; numbers: minutes after release). For all genes except for histones, pre-mRNA levels are shown.

**Supplementary Information, Fig.S 2.**
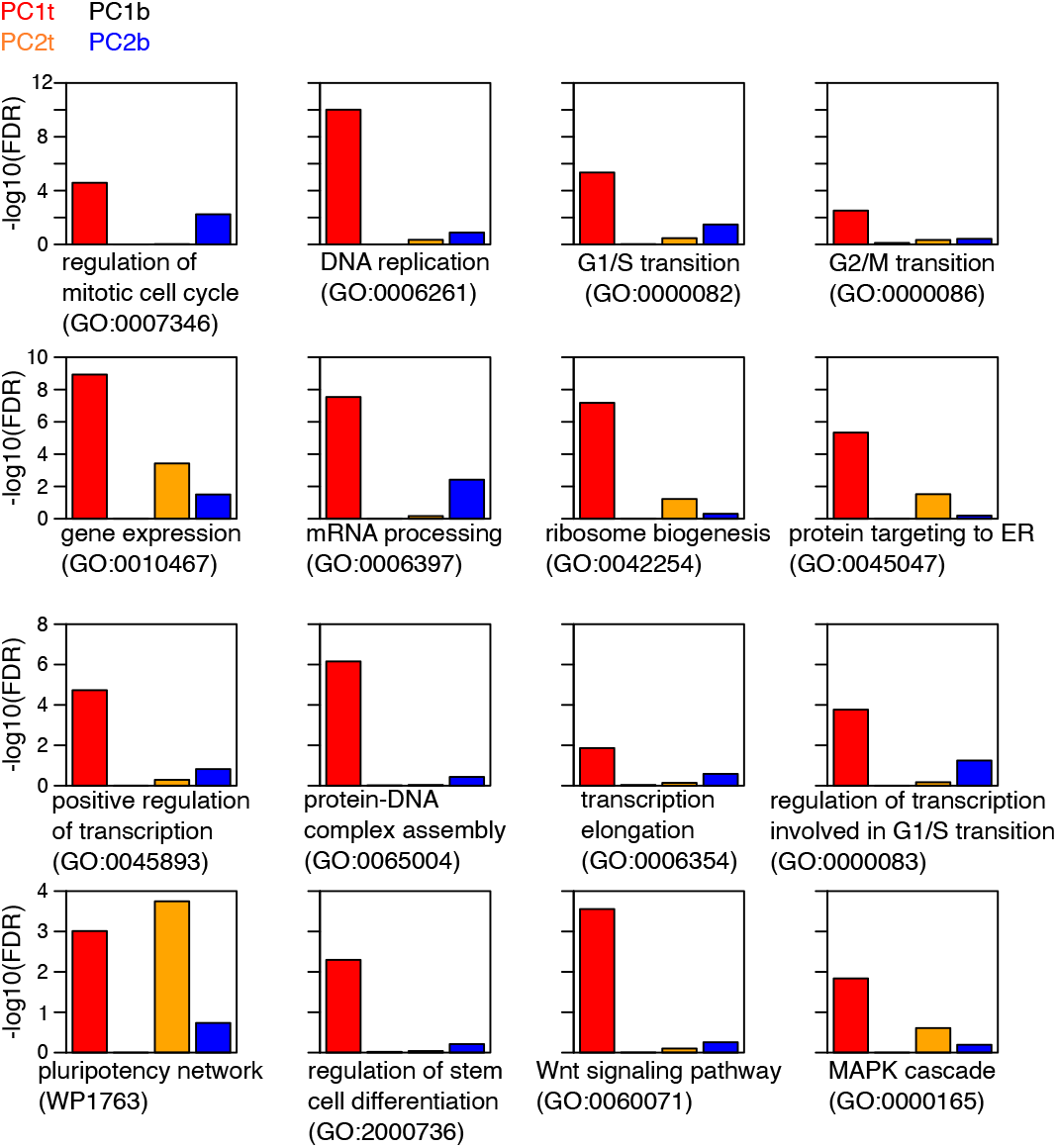
Examples of gene ontology associations on classes of genes showing different post-mitotic behaviour. Gene ontology enrichments (-log10(FDR)) for cell-cycle categories (first row), gene regulation (second and third rows) and stem cell related terms (forth row), calculated for 4 lists of genes derived from PCA analysis of post-mitotic gene reactivation datasets. PC1t, PC1b, PC2t, PC2b correspond to the gene groups shown in Figure 2.

**Supplementary Information, Fig.S 3.**
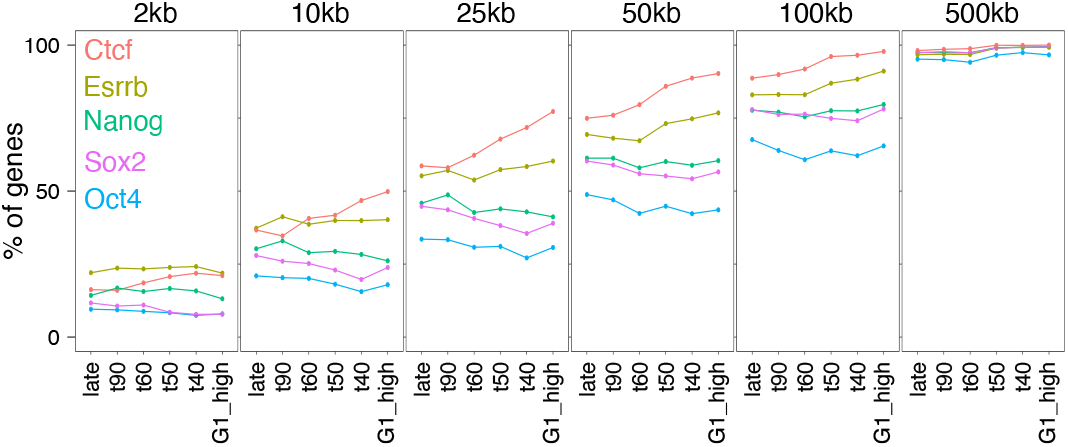
Binding of TFs at increasing distances from gene promoters. Percentage of genes presenting at least 1 TF binding site as detected by ChIP-seq within increasingly bigger genomic windows centred on the TSS, as indicated, calculated for each gene category as in Fig. 3C.

**Supplementary Information, Fig.S 4.**
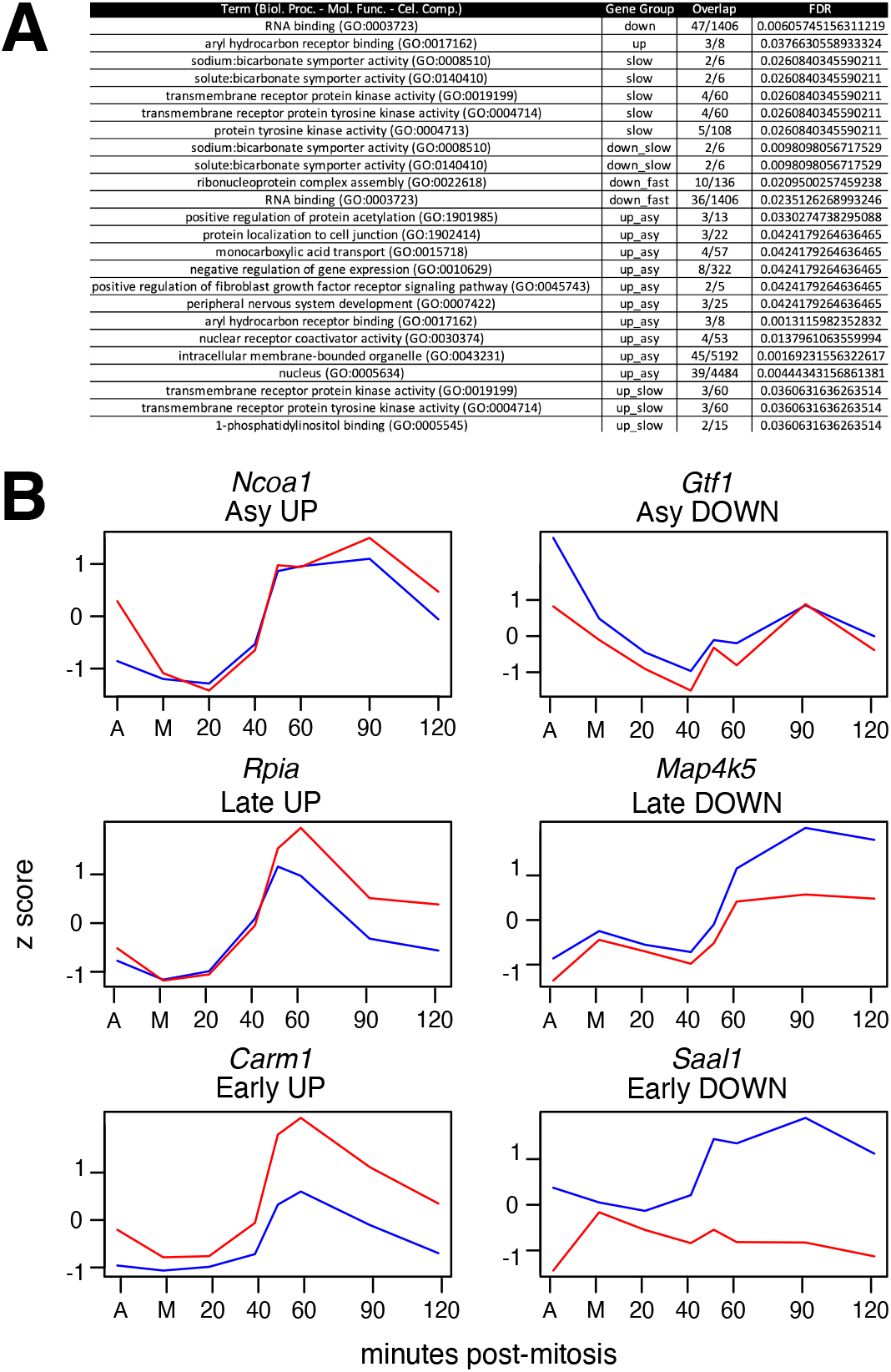
Functional gene groups and examples of pre-mRNAs responding to CTCF depletion. **(A)** Table of all gene ontology terms (FDR<0.05) identified when considering different combinations of genes derived from Fig.5A, as indicated (Gene Group). **(B)** Examples of individual genes belonging to the CTCF responsive gene groups identified in Fig.5A.

